# Diversity at the *HYP1* locus in potato cyst nematodes does not result from developmentally-programmed somatic mutations

**DOI:** 10.64898/2026.07.21.739894

**Authors:** Vincent C. T. Hanlon, Unnati Sonawala, Cian A. A. Raza, Luisa Kalkert, George Harpum, Lukas A. Burkhardt, Johannes Helder, Sebastian Eves-van den Akker

**Affiliations:** Crop Science Centre, Department of Plant Sciences, University of Cambridge, Cambridge, UK; Institute of Microbiology, Department of Biology, ETH Zürich, Zürich, Switzerland; Laboratory of Nematology, Department of Plant Sciences, Wageningen University, Wageningen, The Netherlands

## Abstract

Most genetic diversity stems from spontaneous mutations, that is, errors in DNA repair or replication. But for dozens of organisms across the tree of life, mutations at specific loci are not spontaneous but developmentally programmed: effectively, some organisms edit their own DNA sequences. This is perhaps most common among pathogens and parasites, many of which use editing to diversify genes that produce important antigens. Plant-parasitic potato cyst nematodes are damaging agricultural pests that establish a lifelong feeding site inside the root of their host plant. We previously observed extensive diversity of rare alleles at *HYP1*, the most highly expressed gene that encodes a protein secreted by potato cyst nematodes during parasitism. Importantly, *HYP1* alleles differ from each other by complex, in-frame rearrangements of short repeated sequence motifs within a single exon. Combining several lines of evidence, we previously hypothesized that potato cyst nematodes use developmentally-programmed mutations, or editing, to diversify *HYP1* alleles in the soma. In the current work, we now test this hypothesis. We employ highly accurate long-read DNA sequencing of a simplified genetic system to identify potential rare edited alleles, we use a transgenic yeast system to describe large *de novo* mutations at *HYP1*, and we interpret our findings in light of key population genetic parameters as well as the genetic diversity surrounding *HYP1* and across the genome.

**Significance Statement:** In parasites, genetic diversity is valuable fuel for the coevolutionary arms race with their hosts. However, in parasitic nematode worms, low genetic diversity is often observed. Several other parasites and pathogens (e.g., single-celled eukaryotes) generate genetic diversity in an unusual way: they edit the DNA sequence of an important gene in their own genome, instead of waiting for rare spontaneous mutations to occur. This has never been observed in a plant parasite, but we previously described a major parasitism gene (*HYP1*) in plant-parasitic nematodes with such diverse DNA sequences that it looks like it could perhaps be edited. Now, we generate better data for a simplified genetic system and show that much of the observed *HYP1* diversity was actually cryptic DNA sequencing errors. Although there is real genetic diversity at *HYP1*, it is best explained not by editing but by fundamental evolutionary forces that operate across the genome.

## Introduction

Many parasitic nematode genomes harbour little genetic diversity (Cole & Viney 2018; Koutsovoulos et al. 2020). This is in stark contrast to their free-living relatives, particularly outcrossing species, which often have extremely high levels of genetic diversity (Cutter et al. 2013). Such patterns can be explained by population genetics theory, which predicts that selectively neutral genetic diversity increases with the effective population size at equilibrium (Nei & Li 1979). Whereas free-living nematodes typically have large census population sizes and efficient dispersal (which increase effective population size), the profound dependence of parasitic nematodes on their host plant or animal can often limit dispersal and promote inbreeding, lowering effective population sizes, strengthening genetic drift, and accelerating the loss of genetic variation genome-wide (Cole & Viney 2018).

Conversely, a broad range of organisms—particularly pathogens and parasites—use developmentally-programmed mutations, or “editing”, to locally increase genetic diversity at targeted loci in their own genome (Hanlon et al. 2025). Many pathogens edit the DNA sequences of genes that produce key antigens to escape recognition by the host immune system. For example, the bacteria, fungi, and protozoa that cause syphilis, sleeping sickness, and a form of pneumonia in humans are all believed to perform such editing (Hanlon et al. 2025). Likewise, extensive somatic editing diversifies antigen receptor genes in the immune systems of jawed vertebrates (Tonegawa 1983) and, separately, of jawless vertebrates (Pancer et al. 2004). However, no such editing has yet been confirmed among the parasites or pathogens of plants.

We recently hypothesized that the allelic diversity observed at an important parasitism gene in potato cyst nematodes (*Globodera pallida* and *G. rostochiensis*; Fig. S1) is the product of developmentally-programmed mutations in the soma (Sonawala, Beasley, et al. 2024), that is, somatic editing. The gene in question, *HYP1*, is required for successful parasitism of the host potato plants, during which it is the most highly expressed *G. pallida* gene encoding a secreted protein (Eves-van den Akker et al. 2014). *HYP1* contains striking sequence diversity within the second exon: approximately 8 distinct sequence motifs (15 or 18 bp long) are repeated in complex permutations that form a repetitive domain of ∼400 bp. Using long-read Oxford Nanopore Technologies sequencing (“ONT”) of native DNA and PacBio sequencing of amplicons, we previously observed that (i) *HYP1* alleles differ by complex, in-frame rearrangements of this repetitive domain, (ii) sequence reads from a single population contain many *HYP1* alleles, including many rare alleles, and (iii) PCR of *HYP1* from individual diploid worms yields more than two alleles (Sonawala, Beasley, et al. 2024). To explain these results, we hypothesized that much like the pathogens of humans described above, *Globodera* nematodes might edit the DNA sequence of *HYP1* in some somatic cells, rearranging sequence motifs within the repetitive domain to diversify *HYP1* proteins and promote parasitism (Fig. 1).

**Figure 1.**
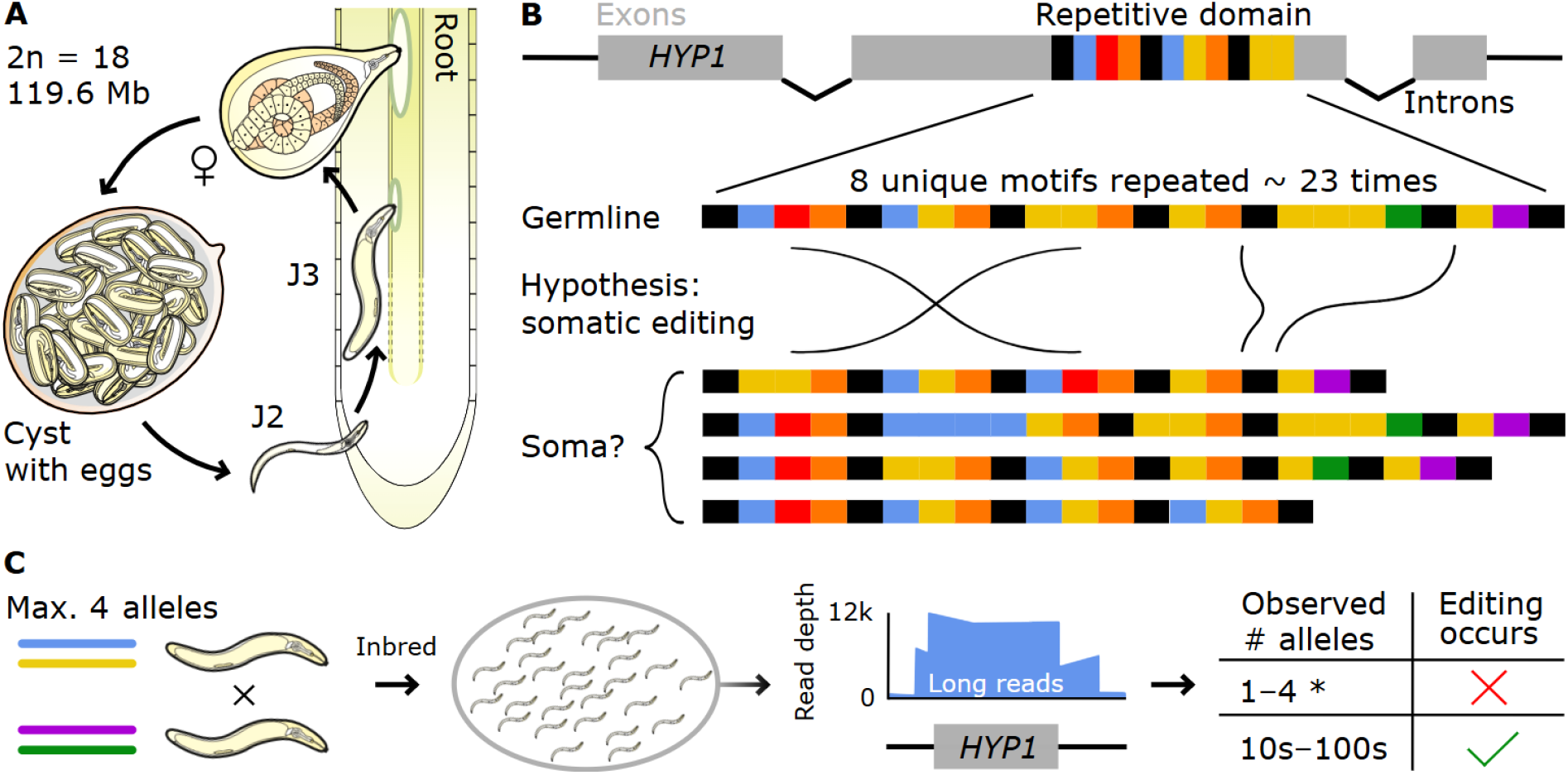
Experimental overview. A. Simplified *Globodera* lifecycle with ploidy and genome size. J2, J3: stage 2, 3 juvenile. Males not shown. B., Schematic showing how the somatic editing hypothesis could explain allelic diversity at *HYP1*. C., Experimental approach for the *G. rostochiensis* inbred population derived from just 2 worms. Sharp changes in read depth around *GrHYP1* reflect the 4 Cas9 guide RNA cut sites. * Additional very rare alleles may also arise by spontaneous (non-developmentally-programmed) mutations.

In this Letter, we test the hypothesis that the observed allelic diversity at *HYP1* results from somatic editing, and we interpret our findings in the context of estimated effective population sizes as well as genetic diversity at the broader *HYP1* locus and across the genome.

## Results and Discussion

To distinguish rare edited somatic alleles from inherited germline variation, we examined an inbred population that descends exclusively from a single mating event (Janssen et al. 1990), in which at most 4 germline *HYP1* alleles can have been inherited from the two diploid *G. rostochiensis* ancestors (“*GrHYP1*”; Fig. 1C). This allowed us to design an unequivocal test of the editing hypothesis: somatic editing occurs only if pooled sequencing of unamplified DNA from this inbred population reveals tens or hundreds of rare *GrHYP1* alleles beyond the 4 that could have been inherited.

We obtained 12,268 reads covering the putatively edited repetitive domain within *GrHYP1* using Cas9-enriched long read ONT sequencing of native DNA. Most reads had reliable base quality scores at *GrHYP1*, with a mean of 28.6 (i.e., 0.14% error probability; SD 6.8; Fig. S2).

We used a reference-free approach to identify reads that contained *GrHYP1*, locate the repetitive domain, and parse each read into an interpretable *GrHYP1* allele consisting of a string of sequence motifs (15 or 18 bp long; see Methods). Two common alleles accounted for 11,378 of the 11,417 reads that could be parsed in this way, with 39 reads apparently supporting rare alleles.

Under the somatic editing hypothesis, rare edited alleles should have large, complex rearrangements of sequence motifs relative to common germline alleles. We manually examined the 39 ONT reads bearing putative rare alleles by plotting the best alignment of each read against one of the two common alleles, coloured by base quality. This revealed that 37/39 reads were likely sequencing errors that differed from a common allele only by small variants embedded in lower-quality sequenced bases. The 2 remaining reads both had insertions embedded in good base qualities (33 and 132 bp; mean phred 29.8 and 36.6 for the repetitive domain), and may represent rare inherited germline variants, spontaneous mutations, or cryptic sequencing errors (Fig. 2; Fig. S3). Given that we found just 2-4 *GrHYP1* alleles among the 11,378 sequences from the inbred *G. rostochiensis* population, we concluded *GrHYP1* is not diversified by somatic editing.

**Figure 2.**
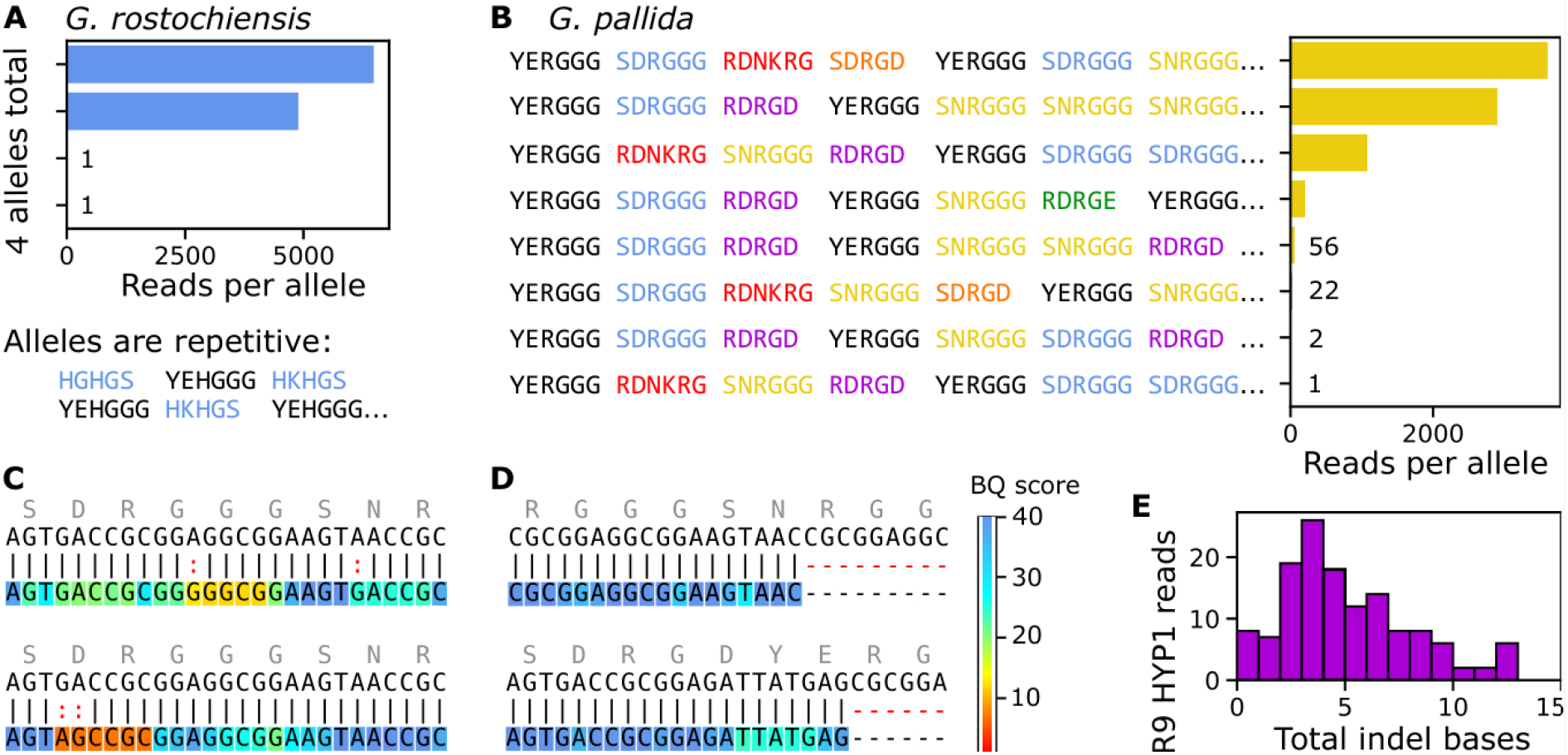
Observed *HYP1* repetitive domain alleles. A. We see 2 common alleles and 2 putative rare alleles in the inbred population (Table S1). *G. rostochiensis* alleles have a simple tandem repeat structure. B. In a non-inbred *G. pallida* population we see 8 alleles total, which consist of complex rearrangements of sequence motifs (Table S2). C. Sequencing errors. For two putative rare alleles (bottom tracks; *G. pallida* R10 data), we show a portion of the alignment against the most similar ground-truth allele (top tracks). D. Two putative rare alleles that passed manual examination, indicated by large indels embedded in good base quality (BQ) scores. E. The old R9 reads match the 6 ground-truth *G. pallida* alleles, as shown by the cumulative length of all indels in alignments between each R9 read and the most similar ground-truth allele at the repetitive domain. Any edited alleles should contain rearrangements of 15 or 18 bp sequence motifs, and should therefore have 15 bp of indels at minimum (Fig. S5).

In our previous work, *G. pallida HYP1* alleles (“*GpHYP1*”) were more diverse and more complex than *G. rostochiensis* (Sonawala, Beasley, et al. 2024). We therefore decided to re-examine *GpHYP1* diversity in the same non-inbred *G. pallida* population studied previously, recognizing that ONT sequencing accuracy has improved substantially since the original publication (R9 vs R10 flow cell chemistry). After performing the same target-enriched sequencing and reference-free analysis as above, we obtained 7954 *G. pallida* reads with parsed *GpHYP1* alleles, including 3 common alleles supported by >1000 reads, 3 intermediate-frequency alleles supported by >20 reads, and at most 2 manually-validated rare alleles (Fig. 2B-D; Fig. S4). The top 6 alleles were the same as those found in the 2024 study. However, our updated sequencing dataset contained far fewer rare alleles than previously observed, suggesting that developmentally-programmed somatic editing does not additionally diversify *GpHYP1* either.

To assess whether this reduced *GpHYP1* diversity might result from an unknown confounder, such as differences in growth conditions, we jointly analyzed the old R9 chemistry ONT reads from our previous work (Sonawala, Beasley, et al. 2024) with ground truth allele sequences derived from our greatly improved R10 dataset. This showed that every old ONT *GpHYP1* read closely matched one of the top 6 alleles in the updated dataset, with at most 12 bp of indels flanked by low base qualities, indicating sequencing errors (Fig. 2E; Fig. S5). It may be that the structure of the *GpHYP1* repetitive domain somehow impairs ONT sequencing: even in the high-quality R10 dataset, we observed 15 bp deletion errors in 13 of the 7954 parsed reads, all in DNA sequences corresponding to “DRGDYERGGG” (Fig. S6). We additionally attribute the *GpHYP1* diversity previously observed in single-worm amplicon sequencing (Sonawala, Beasley, et al. 2024) to ubiquitous PCR template switching errors, which coincided with those observed from ONT sequencing and the hypothesized pattern of edited rearrangements.

Clearly, *HYP1* diversity consists of germline alleles that arise by ordinary mutational processes rather than developmentally-programmed somatic editing. What mutations produce the complex sequence motif structure that defines *HYP1*? To address this question, we constructed a transgenic yeast system to enrich and sequence yeast colonies with mutated *GpHYP1* (Fig. 3). We obtained 76 ONT reads representing *GpHYP1* alleles sampled from ∼120 pooled yeast colonies. Of these, 2 contained probable gene conversion tracts, 2 had an unaltered *GpHYP1* sequence, and 72 contained large deletions. At least 67 mutations maintained the *GpHYP1* motif structure, consistent with homology-directed rearrangements (Fig. 3B-D; Supplementary Material). It seems plausible that erroneous homologous DNA repair of double-strand breaks, as reported for a similar yeast system (Dalin et al. 2025), is the dominant mutational mechanism for the *GpHYP1* repetitive domain.

**Figure 3.**
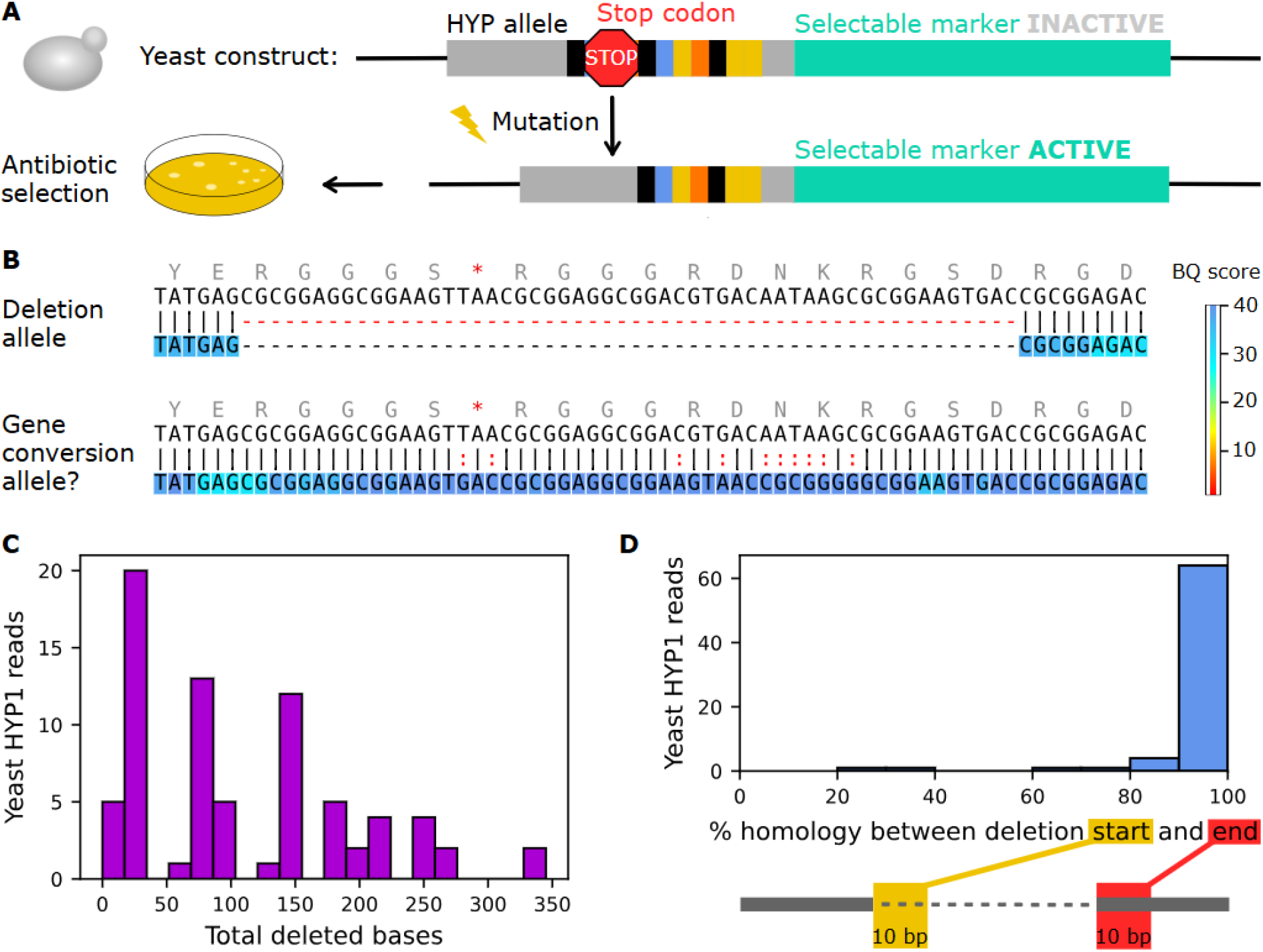
Mutations of the *GpHYP1* repetitive domain in transgenic yeast. A. Simplified schematic of the experimental setup. B. Partial alignments of two sequenced *GpHYP1* alleles in yeast (bottom tracks) against the allele from the original construct (top tracks), coloured by base quality (BQ). In the gene-conversion-like allele, DNA sequence corresponding to S*RGGGRDNKRG has been replaced with SDRGGGSNRGGG from downstream. Stop codons denoted “*”. C. For each read, the total number of deleted bases relative to the original repetitive domain sequence. Most reads were dominated by one large deletion spanning the stop codon. D. Nearly all deletions were embedded in sequence homology. Homology: fraction of matching bases. Only the largest deletion in each read (>10 bp) is considered.

The genomic region surrounding *GpHYP1* contains a striking diversity of single-nucleotide variants (SNVs) and large insertions/deletions within a single population (Fig. 4). The genomes of some nematode species are punctuated by “hyperdivergent regions” with dramatically elevated genetic diversity, which are often enriched for genes related to environmental or pathogen response (Lee et al. 2021; Moya et al. 2025). Investigating this possibility, we began by constructing ONT-based local assemblies for the haplotypes corresponding to the top 4 *GpHYP1* alleles (lengths 57.5-92.3 kb). Then, we combined the four haplotypes into a local graph assembly (see Methods). Examining this graph assembly revealed that four haplotypes are indeed interrupted by large, complex insertions/deletions (35 variants >1 kb) and are broadly collinear. Aligning the ONT reads to the graph assembly allowed us to call SNVs and estimate the mean nucleotide diversity at *GpHYP1* in our population based on the frequencies of the four haplotypes: this was 0.0137, corresponding to 1 SNV every 16 bp.

**Figure 4.**
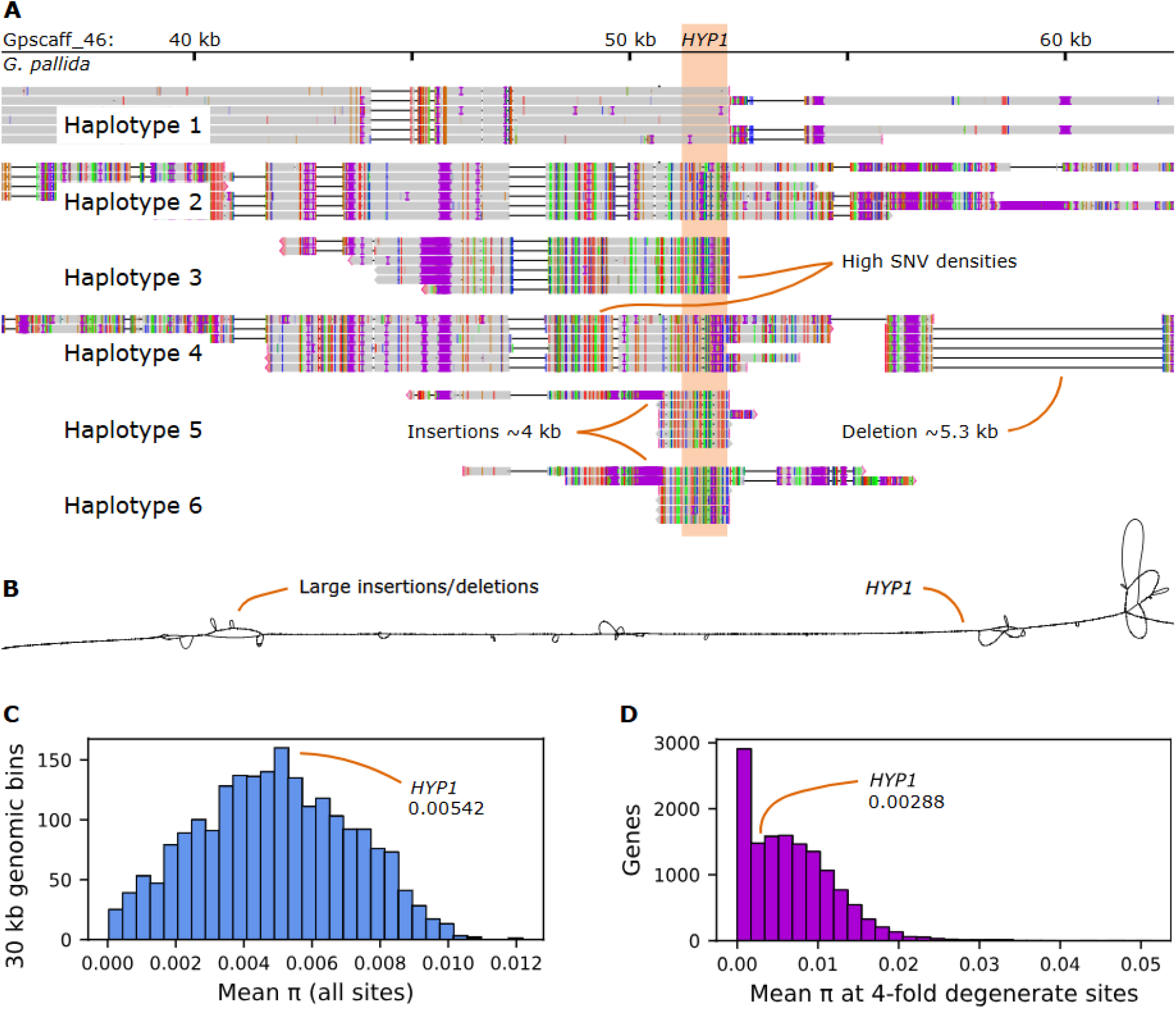
Genetic diversity of *GpHYP1*, the local region, and the genome. A. Diversity among haplotypes near *GpHYP1*. In this genome browser snapshot, ONT reads from a population are grouped by the *GpHYP1* allele they bear. The sharp dropoff in coverage near *GpHYP1* is a consequence of Cas9 enrichment. Note that many reads align only partially because of reference bias. B. Reference-free graph assembly of the top 4 haplotypes (1-4 above) for a region corresponding to approximately Gpscaff_46: 40-65 kb on the linear reference genome (as in A). C. Mean nucleotide diversity (π) calculated on 30 kb bins using *G. pallida* short-read data on the linear reference genome (low-coverage genomic regions removed). Note that π here is approximately 2-fold lower than estimates from ONT long reads on the local graph assembly, perhaps due to reference bias. D. Mean nucleotide diversity (π) calculated at 4-fold degenerate sites, binned by gene. π is zero-inflated, largely because short genes may contain few 4-fold degenerate sites.

Although this is similar to hyperdivergent regions in species such as *Caenorhabditis briggsae* (Moya et al. 2025), hyperdivergent regions ultimately must have higher nucleotide diversity relative to the rest of the genome. We therefore aligned pre-existing short read DNA sequence data for a similar *G. pallida* (Newton) population (Cotton et al. 2014) to the linear reference genome assembly (Sonawala, Beasley, et al. 2024), called SNVs, and calculated the mean nucleotide diversity over 30 kb bins (Fig. 4C-D; Materials and Methods; Supplementary Material). Far from being unusual, this showed that the nucleotide diversity surrounding *GpHYP1* is typical of the *G. pallida* genome, suggesting that it is best explained by genome-wide neutral processes rather than local hyperdivergence.

For populations at equilibrium, high neutral nucleotide diversity corresponds to large effective population sizes (Nei & Li 1979). At four-fold degenerate sites, which are thought to evolve nearly neutrally, the mean nucleotide diversity genome-wide (π) in our *G. pallida* (Newton) population is 0.00746. Given the relationship π = 4N_e_μ and using 1.84 × 10^-9^ from the model nematode *C. elegans* for the mutation rate μ (Konrad et al. 2019), we would expect an effective population size (N_e_) of approximately 1,000,000. This contrasts sharply with reported N_e_ for both *G. pallida* and the related beet cyst nematode *Heterodera schachtii*, which is 64 and 85, respectively (Jan et al. 2016; Montarry et al. 2019).

We can learn something about cyst nematode populations from this discrepancy of four orders of magnitude in the estimated N_e_. The relationship π = 4N_e_μ can produce inflated estimates of N_e_ with even small amounts of migration (Waples 2025). Moreover, it corresponds to the long-term, metapopulation N_e_—whereas the smaller N_e_ estimates are short-term and local, reflecting fundamental aspects of *G. pallida* life history such as inbreeding (Montarry et al. 2019). The picture that emerges is that N_e_ may be low (i.e., strong genetic drift) in small *G. pallida* subpopulations restricted to a particular plant or patch, but that migration between subpopulations may maintain relatively high genetic diversity and perhaps also increase overall metapopulation N_e_. Because N_e_ determines the efficacy of natural selection, it is of great practical importance for assessing how quickly virulent nematode strains are likely to arise that overcome genetic resistance in commercial potato varieties.

In sum, we find that allelic diversity at the parasitism gene *HYP1* does not arise by somatic editing as hypothesized previously. Most likely, the mutations that produce *HYP1* alleles over evolutionary time are homology-directed rearrangements of sequence motifs. We describe elevated rates of nucleotide diversity at *GpHYP1* and across the *G. pallida* genome, perhaps maintained by migration despite small local effective population sizes. This work highlights the impact of recent progress in the accuracy of major DNA sequencing technologies, as well as the importance—and the challenges—of interrogating the genomes of non-model organisms.

## Materials and Methods

### Growth conditions and sequencing

Inbred “Line 19” *G. rostochiensis* and non-inbred *G. pallida* (Newton) nematodes were grown separately on Desiree potatoes in soil. Cysts were extracted by the Fenwick can method and cleaned by acetone floatation (van Bezooijen 2006). Cysts were either crushed between an aluminium block and a glass slide then floated on 50% iodixanol to obtain *G. rostochiensis* eggs (Supplementary Material), or hatched in tomato root diffusate to obtain *G. pallida* J2s. We used phenol-chloroform to extract DNA and performed Cas9 enrichment as described previously (Sonawala, Derevnina, et al. 2024), followed by ONT sequencing on the promethION R10 flow cell (Ligation sequencing DNA v14; Table S3; Supplementary Material).

### Reference-free analysis of alleles

We selected all reads that contained a perfect match for either a known *HYP1* sequence motif or one of 6 arbitrarily-chosen 18mers from *HYP1* exon 2. We parsed the reads by replacing any DNA sequence that matched a known *HYP1* motif (up to edit distance 1) with its amino acid sequence. We discarded reads for which the repetitive domain could not be completely substituted into amino acid sequences in this way. *HYP1* alleles were considered identical if they were parsed into the same amino acid sequence, which allowed us to count the reads supporting each unique allele in a population.

For re-analysis of the R9 ONT reads from the 2024 study (Sonawala, Beasley, et al. 2024), we manually examined all 253 *GpHYP1-*containing reads with mean base quality greater than 16 (as in Fig. S3).

### Transgenic yeast

We assembled the TEF1 promoter, *GpHYP1* exon 2 interrupted by a stop codon, a rigid linker (“PAPAP”), and the antibiotic resistance gene AUR1-C into the pAbAi backbone (Takara) using a combination of overlap PCR, NEBuilder HiFi (New England Biolabs, “NEB”), and mutagenesis PCR, linearized it at *URA3* with BbsI-HF (NEB) and integrated it into the Y1Hgold yeast genome by lithium acetate heat shock (Gietz & Schiestl 2007). We plated the yeast with 200 ng/mL Aureobasidin A (Supplementary Material), scraped off all colonies after 3 days, washed twice with 0.9% NaCl and 0.01% TWEEN-20, and resuspended them in DNA/RNAshield (Zymo Research) for genomic DNA extraction and PCR-free ONT sequencing (Plasmidsaurus).

### Graph assembly and variant analysis

Using *GpHYP1* ONT reads larger than 20 kb, we constructed allele-specific local assemblies either using flye v2.9.5-b1801 with --no-alt-contigs (Kolmogorov et al. 2019) or simply by choosing the two reads with greatest genomic span, aligning them to each other, and taking the consensus sequence. We polished the assemblies with racon v1.5.0 (github.com/isovic/racon) and then constructed a combined graph assembly with pggb v5134732 (Garrison et al. 2024). We aligned ONT reads to the graph using GraphAligner v1.0.17 (Rautiainen & Marschall 2020) and called SNVs relative to the most common *GpHYP1* allele using vg v1.71.0 (Garrison et al. 2018). We aligned short reads (ENA accession ERR123954) to the linear reference assembly using NextGenMap v0.5.5 (Sedlazeck et al. 2013), called SNVs with freebayes v1.3.8 (Garrison & Marth 2012), and identified 4-fold degenerate sites with degenotate v1.2.4 (Mirchandani et al. 2024). Both variant callsets included SNVs (with QUAL>20) and invariant sites (Brault et al. 2026), used to estimate nucleotide diversity under the assumption that every read represents a distinct individual in pooled sequence data from thousands of nematodes (Supplementary Material; Figs. S7-S8).

## Acknowledgment

We are grateful to Victor Hugo Moura de Souza, Christopher Stephens, Anika Damm, Madalena Mendonça, Olaf P. Kranse, Priya Desikan, and other members of the Plant-Parasite Interactions Group for training and discussions related to nematology and microbiology techniques. We also thank Matthew Back for providing the dimensions of the cyst crushing apparatus.

## Data Availability

ONT reads for the *G. rostochiensis* inbred population, the *G. pallida* (Newton) non-inbred population, and pooled transgenic yeast colonies are available from the NCBI Sequence Read Archive (BioProject PRJNA1496558). Analysis files are available from the Dryad Digital Repository (doi.org/10.5061/dryad.1c59zw4cd), including *G. pallida* variant calls, gene coordinates, the graph assembly, all alignment PDFs for manual examination, and the .sat file for the cyst crushing apparatus. All analysis scripts can be found on GitHub (github.com/vincent-hanlon/hyp-globodera).

